# Schema Playground: A tool for authoring, extending, and using metadata schemas to improve FAIRness of biomedical data

**DOI:** 10.1101/2021.09.02.458726

**Authors:** Marco Cano, Ginger Tsueng, Xinghua Zhou, Laura D. Hughes, Julia L. Mullen, Jiwen Xin, Andrew I. Su, Chunlei Wu

## Abstract

**Background:** Biomedical researchers are strongly encouraged to make their research outputs more Findable, Accessible, Interoperable, and Reusable (FAIR). While many biomedical research outputs are more readily accessible through open data efforts, finding relevant outputs remains a significant challenge. Schema.org is a metadata vocabulary standardization project that enables web content creators to make their content more FAIR. Leveraging schema.org could benefit biomedical research resource providers, but it can be challenging to apply schema.org standards to biomedical research outputs. We created an online browser-based tool that empowers researchers and repository developers to utilize schema.org or other biomedical schema projects.

**Results:** Our browser-based tool includes features which can help address many of the barriers towards schema.org-compliance such as: The ability to easily browse for relevant schema.org classes, the ability to extend and customize a class to be more suitable for biomedical research outputs, the ability to create data validation to ensure adherence of a research output to a customized class, and the ability to register a custom class to our schema registry enabling others to search and re-use it. We demonstrate the use of our tool with the creation of the Outbreak.info schema—a large multi-class schema for harmonizing various COVID-19 related resources.

**Conclusions:** We have created a browser-based tool to empower biomedical research resource providers to leverage schema.org classes to make their research outputs more FAIR.

## Introduction

Funding agencies, international consortia, institutional policies, and publisher requirements have helped promote the adoption of the FAIR (Findability, Accessibility, Interoperability, and Reusability) guiding principles (Wilkinson et al, 2016)(Boeckhout et al, 2018) for biomedical research data sharing to varying degrees of success. While it is now standard to make datasets accessible and potentially reusable via deposition of the dataset in a repository, standardization issues continue to make it challenging for researchers to make datasets findable, interoperable, and reusable. To address these issues, domain experts and data stewards have been inspecting the gap between principle and practice (Koesten et al, 2020), extending (Jauer and Deserno, 2020) and adapting the principles (Holub et al, 2018), creating their own metadata standards (Canham and Ohmann, 2016) and data schemas (Hruby et al, 2016)(Papadiamantis et al, 2020). However a large gap remains between the communities that develop standards and the adoption of these standards by data and resource providers due to issues in communication, education/training, incentives, and the availability of supportive tools (Hollman et al, 2020). For example, the Dublin Core Metadata Initiative (DCMI) provides a metadata ontology (i.e.-a structured vocabulary for classifying and describing metadata): terms and data elements (Dublin Core Metadata Initiative, 2020), two general-use schemas (i.e.-sets of metadata vocabulary used to describe a conceptual entity): core and qualified, and a thorough guide for utilizing their ontology with their model-based framework for creating schemas: the Dublin Core Application Profile (DCAP) guide (Coyle and Baker, 2009). The DCAP guide was intended to empower data providers to mix and match Dublin Core (and other) metadata terms/elements (properties) to create new application profiles (schemas) to suit their needs. While the core (data element) schema has been widely-adopted, the lack of authoring tools to help create more type/concept-specific schemas and the lack of tools for transforming schemas into working formats for consumption and implementation has hampered the adoption and implementation of DCAP (Baker, 2019).

Schema.org is a metadata vocabulary standardization project founded by the major search engine companies such as Google, Microsoft, Yahoo, and Yandex. It is an open source, collaborative initiative that develops metadata standards for improved searchability. Schema.org already includes some biomedically relevant classes (i.e.-conceptual entities) like Datasets and Medical Study, and applying schema.org classes to biomedical research resources would improve interoperability, enabling researchers readily ingest existing resources and to leverage search engine-based solutions (like Google Dataset Search) to find resources of interest. Although there have been some efforts to leverage schema.org to improve findability of scientific research data (Sansone et al, 2017)(Papadiamantis, 2020)(Jones et al, (2021) and many generic repositories (like Figshare and Zenodo) are compliant, schema.org remains largely underutilized by the biomedical research community. Bioschemas is an open and collaborative effort that has been actively promoting the use of schema.org in the life sciences by serving as a hub for researchers to create new biomedically relevant classes with the goal of refining and proposing these classes to schema.org (Gray et al, 2017)(Profiti et al, 2018), and by raising awareness about the usefulness of metadata schemas. The Bioschemas community has also identified the need for easy-to-use tools to help improve public accessibility and participation in the schema development process.

Here, we describe the Data Discovery Engine’s (DDE) Schema Playground, a web-based tool that improves the ease of using any registered schema or schema.org classes. Our tool allows users to easily find and visualize relevant classes from Schema.org (Bioschemas, BioLink, and others), extend them, create json schema validation rules, and save/share the newly created classes for others to reuse. Our tool also includes a framework for building data registries and creating guides for data submission; however, the implementation and integration of these features on our site is restricted to partner organizations. We introduce the features of this tool, review its value to different types of users, demonstrate its application towards the creation of a new schema for COVID-19-related resources, and discuss its adoption by the Bioschemas metadata standardization community.

## Implementation

The Data Discovery Engine’s Schema Playground is a browser-based tool built with Vue.js, Python/Tornado, and the BioThings Software Development Toolkit (https://docs.biothings.io/, a framework for building biomedical APIs). Schemas from schema.org and other consortia/projects are stored and made searchable using MongoDB and Elasticsearch. The code for the Schema Playground can be found at https://github.com/biothings/discovery-app and is free to use under the Apache License 2.0. The schema generated by the DDE are exported as json LD files, following json schema and RDF schema specifications. The COVID-19 outbreak.info resource schemas were developed by comparing metadata properties across multiple type-specific repositories to identify properties in common. For example, metadata from LitCovid/PubMed, BioRxiv/MedRxiv, various journals like JAME, NEJM and others, and the metadata from publications found on Zenodo, Figshare and others were compared in order to identify a suitable schema for COVID-19-related publications. Similarly, protocols from protocols.io and the BioSchemas LabProtocol class were compared to develop a schema for COVID-19-related protocols. Once the desired properties and structure for each class of COVID-19-related resource was identified, the schemas were created by extending existing schema.org classes using the DDE Schema Playground.

## Results

The DDE Schema Playground consists of two standard (and fully-accessible) components and two related, custom (limited-access) components (Figure 1). The standard components improve the ease of use of schemas and classes, while the custom components help communities to reap the benefits of their use. The Schema Editor allows users to import community standard schemas like schema.org and customize them for biomedical purposes. These extended schemas can then be shared in the Schema Registry, which allows users to view the schemas and reuse them. When used in conjunction with Data Portals built with BioThings SDK, The DDE Schema Playground can automatically generate data submission forms known as Data Guides.

**Figure 1.**
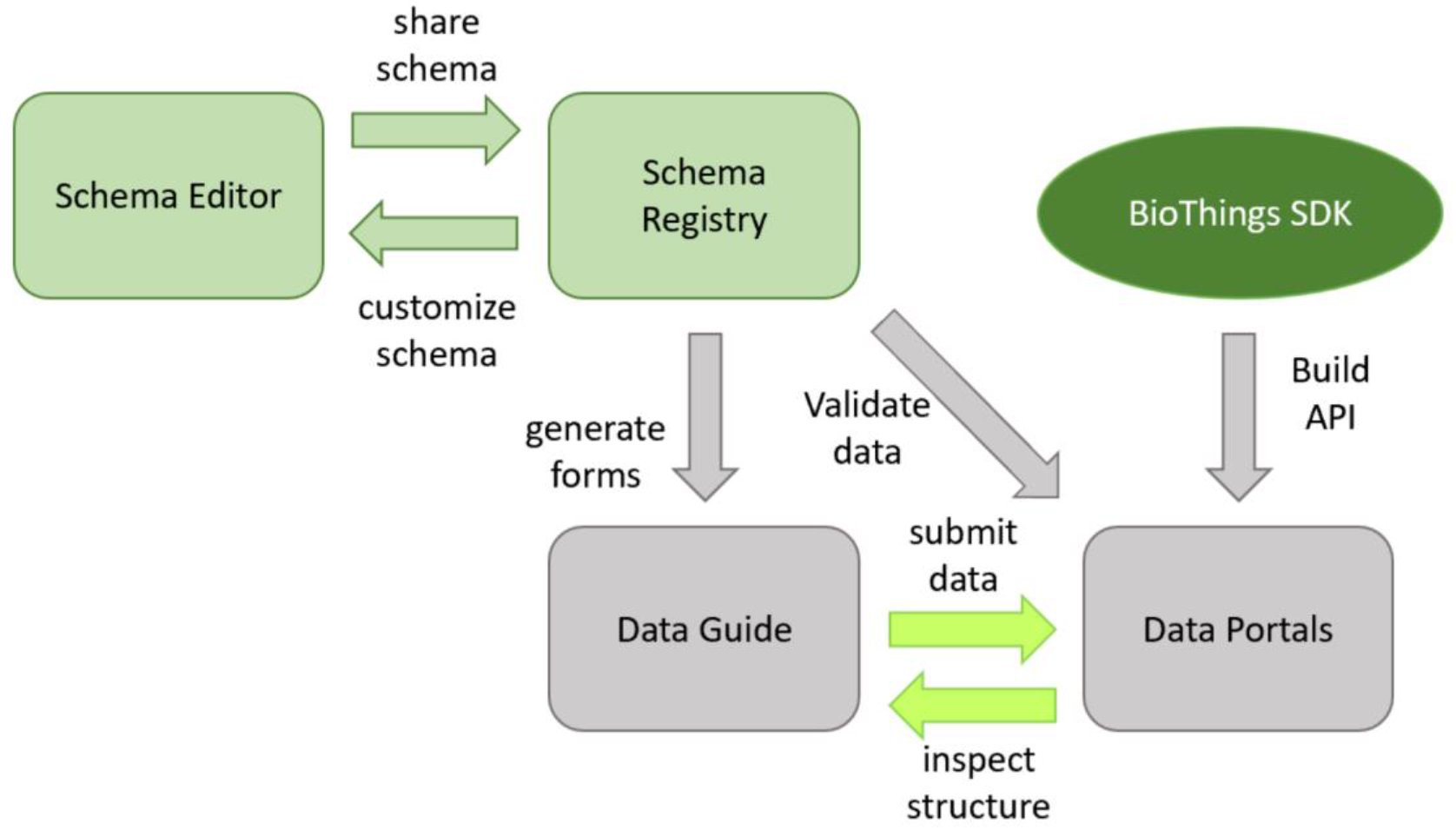
Components of the DDE Schema Playground and how they work together.

To understand how the Schema Playground might help to bridge the gap between data standardization communities and data resource providers, we identified potential utility and value of each of the DDE Schema Playground components for different types of users in our partner communities (Figure 2).

**Figure 2.**
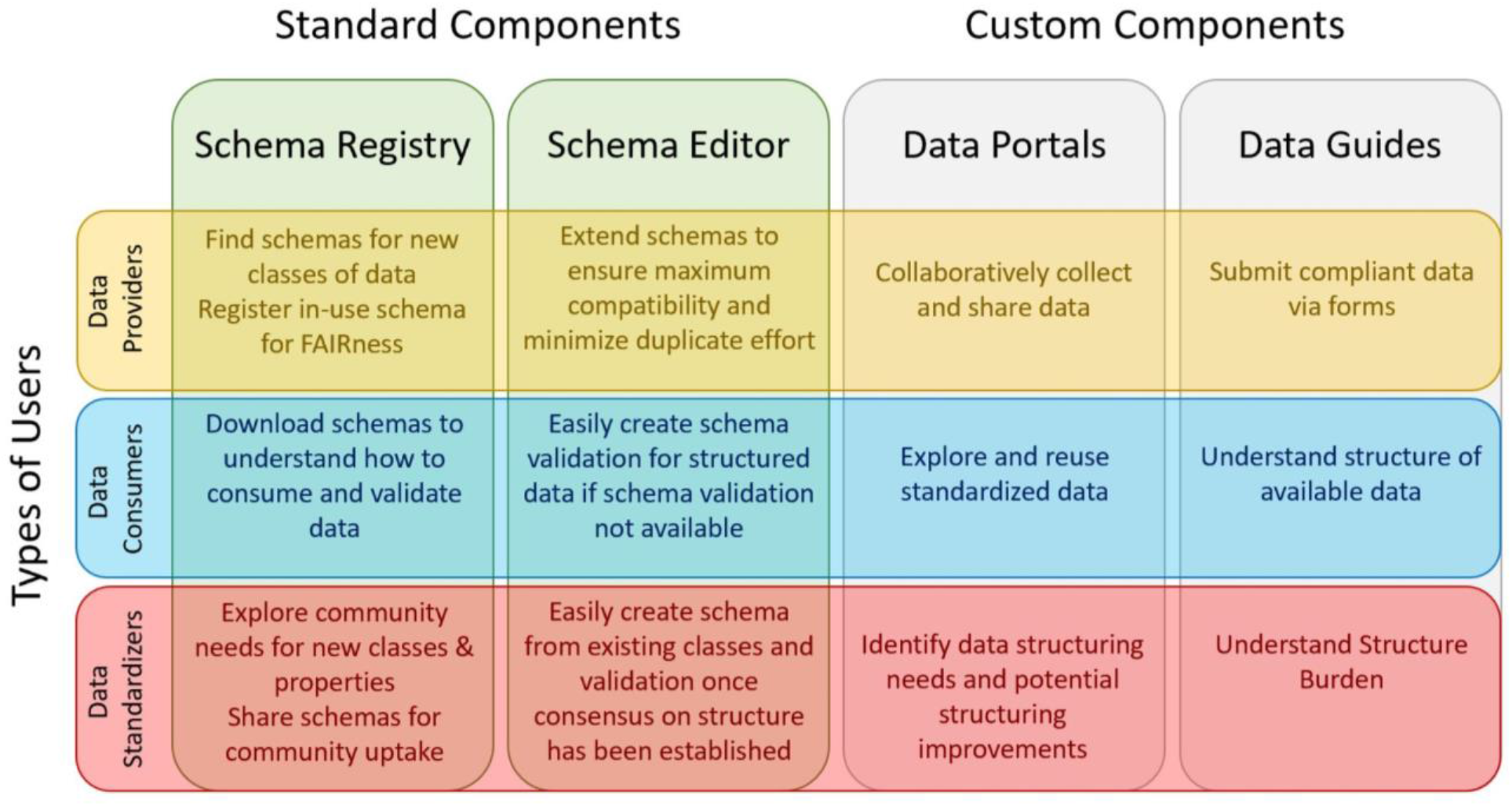
The value and utility of DDE Schema Playground components to different types of users.

Any data portals and guides can be used by anyone with sufficient access rights, but the creation of a data portal or data guide requires partnerships with our team to actualize. For the outbreak portal, data submission via the guide is open to all and utilizes github for authentication. For other portals, access may be restricted as required by the responsible partner organization. The data portal and data guides allow data providers and data consumers to collect, share, and use data. Since the data guide converts a custom schema into a web-based data submission form, it enables data consumers and data standardizers to visually inspect and understand the burden of structure.

The schema registry and editor allows data providers and/or standardizers to find, customize, and share schemas. Sharing schemas via the registry will make it easier for data consumers to understand how to consume data from a data provider and to create data validation if one is not available from the data provider. For example, data-use restrictions usually require a data consumer to create an account with the data provider in order to access data. However, data consumers cannot easily determine whether the data locked behind the account-creation process will actually be useful prior to creating an account. Sharing the data schema via the DDE registry could address this issue by allowing data consumers to understand what’s available without actually displaying any restricted-access data. Having a central location for schemas submitted by data providers will also make it easier for data standardization communities to evaluate the needs of the biomedical research community. To further illustrate the value of the schema registry and editor, we compare and detail the features of the DDE Schema Playground with available tools for creating, applying, and consuming other major schemas such as Schema.org and Bioschemas.

Schema.org, Bioschemas and other data standardization efforts have built strong communities to generate consensus on data modeling for the creation of new schemas or the improvement of existing schemas. Hence, there are extensive processes in place (but few tools) for the creation of a new schema based on schema.org or any other schemas. Because of its widespread adoption, there are third party tools available for utilizing and consuming markup from schema.org. The Bioschemas community has developed a process for defining new classes and has a set of tools which cover both the creation of a new schema (google spreadsheet conversion), utilization of a schema (markup generation), and evaluation of use (markup validation, scraping), but these tools vary in usability based on the users programming experience. In contrast to schema.org, Bioschemas also defines cardinality (allowable number of values per property) and marginality (optional vs required value) in its profile schemas as these are important to the life sciences research community. Although the DDE schema playground was developed independently from the Bioschemas community, our interests aligned and we sought to provide complimentary schema tools to facilitate biomedical schema development and adoption. To do this, we identified schema tools and features available directly from the schema.org and Bioschemas communities. We expanded the list of tools by searching for “schema.org tools”, “schema generation tools”, “schema creation tools”, “schema editing tools”, “schema validation tools”, “bioschemas tools” in google). Most user-friendly tools were aimed towards the generation, extraction, or validation of schema-compatible markup rather than the development of schemas themselves (Supplemental Table 1). The Bioschemas community has a few well-documented tools for schema development, but many of those tools were only available as source code and required basic programming experience. We focused our efforts on features for which user-friendly tools for schema creation and reuse, resulting in a web-based application that empowers individual data resource providers to utilize and customize existing schemas from schema.org and other similar efforts. As seen in Table 1, these features include:

1. **Searching and viewing schemas from schema.org and other metadata standardization efforts** The DDE Schema Playground allows for the visualization of json schemas hosted online either on github or elsewhere (Supplemental Figure 1A). This allows users who are familiar with schema.org to review their compliant schema in a more human readable format. The DDE Schema Playground also has a searchable registry of classes from schema.org, BioLink (Bruskiewich et al, 2021), BioThings, Bioschemas, and others. Users may browse and visualize the schemas for various classes from these sources to identify the classes of most interest to them (Supplemental Figure 1B). If a community like Bioschemas or consortia like N3C is interested in making a new schema available for searching and viewing, they can import and register their json schema. The DDE Schema Playground also enables users to compare up to four schemas. For example, there are multiple Dataset schemas available in the registry, and users can compare them to see what properties are unique to each and what properties they share (Supplemental Figure 1C).
2. **Extending and customizing a pre-existing schema for a particular use** The ability to browse and inspect pre-existing schemas makes it easier for a user to customize or extend the schema to suit their own purpose. All the properties from the pre-existing schema will be inherited in the extended schema; however, the user may select properties for which validation is desirable. The user can also create new properties to be included in the extended schema. For example, the Dataset schema from schema.org serves as a potential foundation, but a schema focused on COVID-19-related datasets may need additional fields (e.g., infectiousAgent). To tailor the Dataset schema, we find and extend it from the registry (Supplemental Figure 2A). After we create a name for our schema (the namespace) and the class, we can customize it. We can select to include any property that is available from the schema we are extending (Supplemental Figure 2B), and we can create new properties (eg-curatedBy) that are tailored to our needs (Supplemental Figure 2C). This feature also serves as an easy way to maintain Bioschemas profiles as users can update a registered profile by extending from it, making the necessary changes, and pushing them back to Bioschemas.
3. **Creating validation for the schema for data quality enforcement** Marginality (whether a property is required or not) and cardinality (whether a property can have one or multiple values) are two aspects of schema properties that are not expressed well by schema.org but are desirable to biomedical researchers (Supplemental Figure 3A and 3C). In the DDE Schema Playground, this is handled via the creation of json schema validation rules, and the DDE’s Schema Validation Editor provides a simple drag and drop mechanism to create straightforward validations (Supplemental Figure 3B). For slightly more complex validations, the user can edit the type of validation they are trying to include before dragging and dropping it into the property of interest. In our example Dataset schema, an Organization is a potential type for our new property (curatedBy). We edit the example object validation for Person to create an Organization object validation (Supplemental Figure 3D).
4. **Exporting and saving a schema generated by the Schema Playground editor** The DDE Schema Playground allows you to export/download your newly created schema locally and it is also integrated with GitHub, allowing users to save to their GitHub repository (Supplemental Figure 4A-C). The integration with GitHub allows the edits to the schema to be made by multiple parties and provides the schema owner the option of pulling changes to the schema. Additionally, the schema can be forked and edited/customized allowing for re-use of the schemas which in turn improves findability and reusability of resources which follow the schemas.
5. **Registering a newly created schema in the DDE schema registry to facilitate its extension and re-use** Once saved in GitHub, users can review their schema with the schema viewer and add it to the registry to enable others to easily re-use it (Supplemental Figure 1A). This provides a user-friendly interface for editing, customizing, and re-using schemas for those who prefer not to manually edit text and format in json.

**Table 1.**
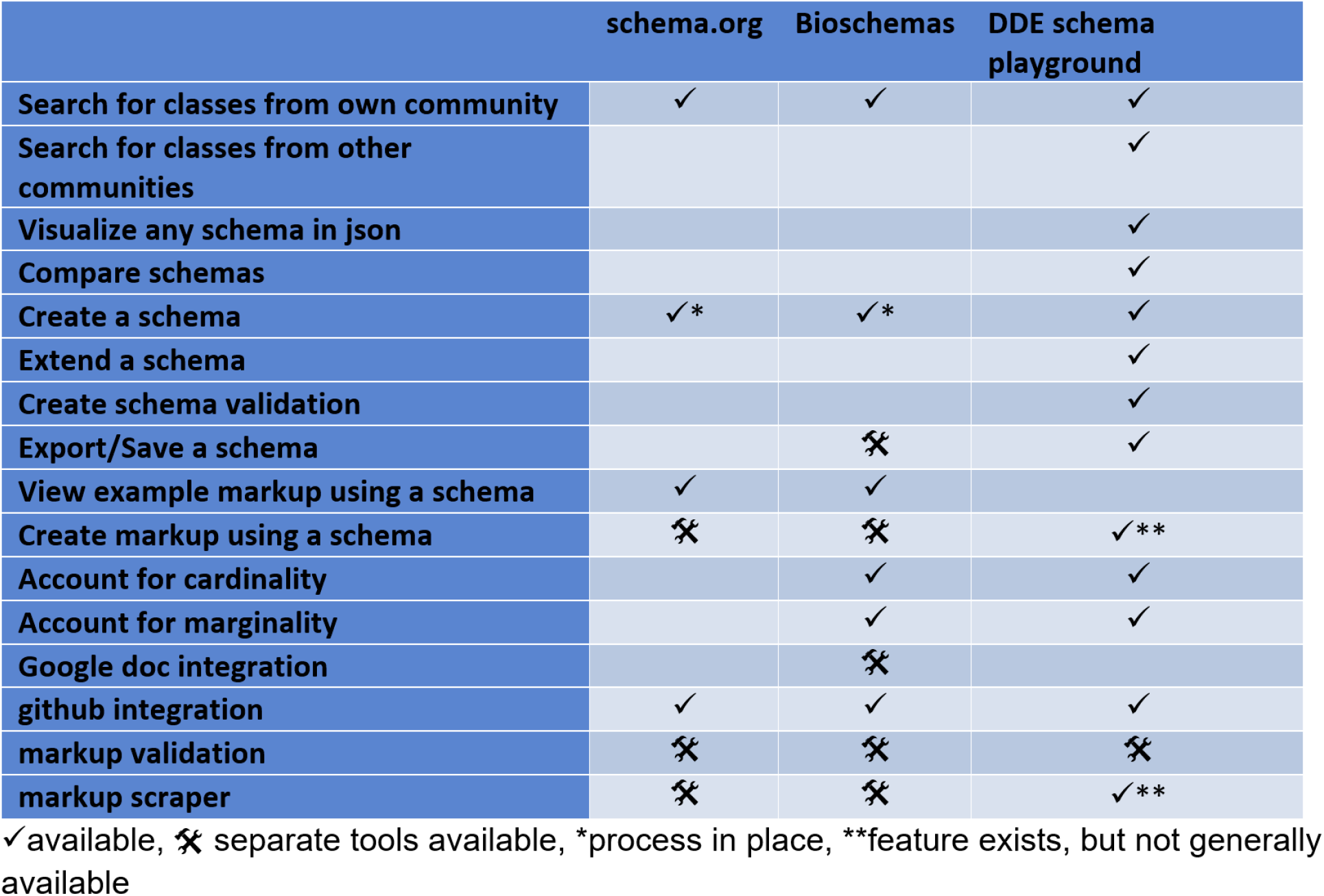
Comparison of Schema.org, Bioschemas, and DDE Schema Playground.

The DDE Schema Playground offers any user the ability to reuse and extend existing schemas. This tool is primarily to assist in the authoring of schemas for use in other applications. In addition, we have converted three Dataset schemas into “guides”, which are web-based forms for annotating resources using schemas authored in the DDE Schema Playground. Annotations created using these guides are stored within a resource registry hosted within the DDE. There are currently three public guides based on the Dataset schemas for the outbreak.info web application (Research Library, 2020), the N3C initiative (Haendel et al., 2020), and the CD2H consortium (Center for Data to Health, 2021). While the creation of guides from schemas is not a fully-automated feature that is available to all users, most of the underlying components are reusable, additional guides can be constructed and hosted within the DDE through collaboration. The Bioschemas community is in the process of integrating the DDE schema playground as part of its schema creation and update process to improve participation by members who lack the programming expertise needed to participate via their previous pipeline.

### Creating the COVID-19 Outbreak schema using the Schema Playground

schema.org classes are often simultaneously too broad (lacking properties needed) and too narrow (including too many irrelevant properties) for a specific research purpose. For this reason, it becomes necessary to adapt schemas to suit needs of a biomedical research project. Outbreak.info is a project from the Su, Wu, and Andersen labs at Scripps Research to unify COVID-19 and SARS-CoV-2 epidemiology and genomic data, published research, and other resources. The standardization of published research and other resources was accomplished by creating a single, multiclass schema to harmonize the metadata: The COVID-19 Outbreak schema. This schema can be found in the DDE registry at https://discovery.biothings.io/view/outbreak/ and was built via the DDE Schema Playground with some manual editing (for merging all the classes into a single schema). There are five principal classes in the Outbreak schema (Analysis, Dataset, ClinicalTrial, Protocol, Publication) and many subclasses to support the principal classes. As seen in Table 2, the classes in the Outbreak schema were extended from related schema.org classes (whenever available) and were created based on metadata comparisons from a variety of related sources. By extending from existing schemas, we reuse existing metadata properties when appropriate, and create new properties only when necessary.

**Table 2.** Classes in the Outbreak schema and how they were created and used.

| Outbreak schema class | extended from schema.org class | inspired by / derived from | used in Outbreak schema class |
| --- | --- | --- | --- |
| Analysis | CreativeWork | Outbreak Protocol, Dataset Class, various COVID-19 Analyses | -- |
| ClinicalTrial | MedicalStudy | NCT Clinical Trials PRS, WHO ClinicalTrials | -- |
| Dataset | Dataset | NIAID schema | -- |
| Protocol | HowTo | Bioschemas LabProtocol, protocols.io | -- |
| Publication | MedicalScholarlyArticle | NLM Medline, ~~bioRxiv~~, various journals | -- |
| ArmGroup | Thing | NCT Clinical Trials PRS, WHO ClinicalTrials | ClinicalTrial |
| CitationObject | CreativeWork | Outbreak primary classes | all |
| Correction | CreativeWork | CitationObject | Publication |
| DataDownload | DataDownload | NIAID schema | Dataset |
| Eligibility | Thing | NCT Clinical Trials PRS, WHO ClinicalTrials | ClinicalTrial |
| Instrument | Product | Bioschemas LabProtocol, protocols.io | Protocol |
| Intervention | Thing | NCT Clinical Trials PRS, WHO ClinicalTrials | ClinicalTrial |
| MonetaryGrant | MonetaryGrant | NIAID schema | all |
| Organization | Organization | NIAID schema | all |
| Outcome | Thing | NCT Clinical Trials PRS, WHO ClinicalTrials | ClinicalTrial |
| Person | Person | NIAID schema | all |
| Product | Product | Bioschemas LabProtocol, protocols.io | Protocol |
| StudyEvent | Thing | NCT Clinical Trials PRS, WHO ClinicalTrials | ClinicalTrial |
| StudyStatus | Thing | NCT Clinical Trials PRS, WHO ClinicalTrials | ClinicalTrial |
| StudyDesign | Thing | NCT Clinical Trials PRS, WHO ClinicalTrials | ClinicalTrial |
| StudyLocation | Place | NCT Clinical Trials PRS, WHO ClinicalTrials | ClinicalTrial |

For example, the level of detail provided by Protocol Registration System (PRS) schema used by the National Clinical Trial (NCT) registry is more granular than schema.org’s MedicalStudy class, but broad enough that it encompasses properties from both child classes of MedicalStudy (MedicalTrial and MedicalObservationalStudy). The child classes of MedicalStudy only differ in the property name for the study design (trialDesign vs studyDesign), and this property is not delineated in PRS. Further, the PRS includes many properties not currently available in any of these schema.org classes. Adopting the PRS directly was also problematic as we planned to ingest records from other registries like the World Health Organization’s Clinical Trial registry (WHOCT), and the PRS was also more granular than WHOCT. For this reason, the Outbreak.info ClinicalTrial class was created by using the DDE to extend from schema.org, leveraging the PRS-WHO crosswalk (WHO/ICJME, 2019), and creating properties that could help with issues previously identified (Miron, Gonçalves, and Musen, 2020).

In addition to adapting schema.org classes to normalize record data from multiple sources within a class, Outbreak.info needed to normalize common metadata properties between different classes. The hierarchical nature of schema.org classes simplified this process, as many derivative classes inherit properties from the Thing class. For example, the Protocol class in the Outbreak schema was extended from the HowTo class in schema.org and was based on properties identified from available metadata in protocols.io and the LabProtocol profile from Bioschemas. Since both the schema.org classes, MedicalStudy and HowTo, are derivatives of Thing, the Outbreak schema naturally has properties in common across multiple classes and can normalize the metadata across these classes allowing for cleaner query design and improved search functionality. This schema is currently used to harmonize and improve FAIRness of metadata from over 200,000 resource entries in the outbreak.info research library at https://outbreak.info/resources.

### Adoption of the Schema Playground into the Bioschemas schema development and maintenance pipeline

Previously, the pipeline for updating a Bioschemas specification involved the use of a google spreadsheet for attaining community consensus, a command-line tool for converting the csv from the spreadsheet to yaml, cloning the Bioschemas website repository and copying/editing html and yaml files, running Jekyll to test the changes, editing example files in the Bioschemas specifications repository, and creating pull requests for the Bioschemas website repository once everything had performed as tested. The level of expertise needed in order to update a specification has been discussed in multiple Bioschemas community calls as a potential barrier to participation. After initial tests during and after Biohackathon 2021, the Bioschemas community has decided to adopt the DDE into its schema development and maintenance pipeline. Manuals for using the DDE to create or update Bioschemas specifications have been developed, and automated scripts using github actions are currently being developed to more tightly integrate the tool into the pipeline. As seen in Figure 3, the process for updating a Bioschemas profile requires less technical expertise after the integration of the DDE. While the process prior to and after the DDE still requires the ability to edit a yaml/json file (brown) and the ability to use GitHub (black), the DDE-based process does not require the user to have the technical knowledge needed to run tools via the command line (green), or to use Jekyll (blue).

**Figure 3.**
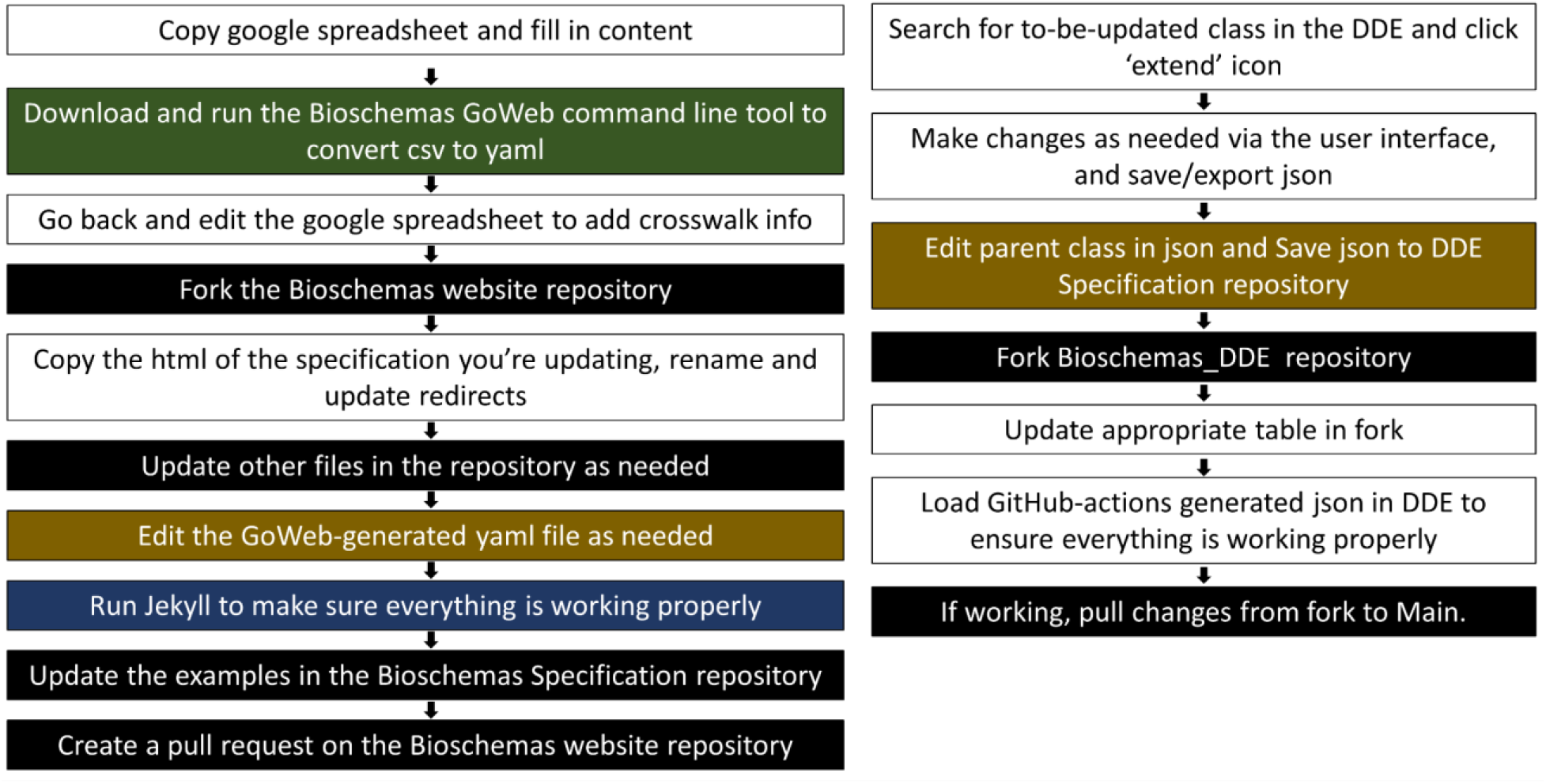
The Bioschemas profile update process before (left) and after (right) the inclusion of the DDE.

## Discussion

In an effort to make scientific resources more FAIR, communities in the biological sciences (Bioschemas), earth sciences (Science on Schema), and more are working diligently to align and influence schema.org to suit the needs of the scientific research community. These communities play an important role in introducing schema.org to the scientific research resource providers and creating tailored schemas more suitable for the research community. Although these communities have helped to create more relevant classes or improve existing classes, it is difficult to push these suggestions to schema.org without compelling use cases or widespread adoption of these tailored classes. For example, the Bioschemas community first introduced the Gene class in 2018, but it was not included as a pending class in schema.org until 2021 due to a lack of widespread adoption. The Bioschemas community spent considerable time and effort on education and training in order to increase the adoption of Bioschemas classes; however, participation in the development of the classes was hampered by the technological expertise needed in order to update a Bioschemas class. The availability of user-friendly tools can make it easier to find and use schema.org and other community-driven schema classes, and empower data providers and researchers to engage in schema authoring and sharing.

Most tools for utilizing existing schema.org classes focus on the utilization of an existing schema (such as markup generation) and lack the ability to customize the schema in a schema-compliant way. Tools that do allow customizing/creating a schema (eg-Bioschemas GoWeb) often require some degree of programming. The DDE Schema Playground is a browser-based tool that enables members of the research community to easily adapt schemas to suit their need and to enable community re-use of their schemas through the DDE schema registry. This encourages and empowers researchers to structure their data in a schema-compliant fashion earlier on in the scientific research process rather than as an afterthought. The schema authoring by the research community, for the research community will encourage the creation and adoption of new classes and properties, which may have previously been neglected due to the absence of representation (ie-expert subject matter volunteers) in data standardization communities. In this fashion, the DDE Schema Playground allows for researchers to express and share their data structuring needs with the data standardization community without diverting attention away from their primary research efforts. Data standardization communities also benefit because their volunteer time can be concentrated on classes already in use by researchers (but could benefit from some standardization), and diverted away from classes which lack interest/support from the research community at large.

The DDE currently only allows the registration of schema (ie-classes described by sets of properties), while many well-used metadata ontologies such as the Web Ontology Language (OWL) or Dublin Core Metadata Initiative (DCMI) exist simply as hierarchies of properties that are not tied to any class. These metadata ontologies intentionally do not group the properties into classes in order to encourage the mix-and-match of properties. Although these non-classed metadata ontologies cannot be registered in their entirety as classless properties in the DDE at this time, the DDE can flexibly ingest properties from any metadata ontology as long as it is properly formatted (ie-conforms to json schema formatting). This means that users can build their schema by extending from schema.org, Bioschemas, or any registered schema, and incorporate properties from OWL, DCMI, and others as needed. For example, all Bioschemas profile classes also include the ‘conformsTo’ property from DCMI, and the NIAID Dataset schema also leverage properties from OWL. In theory, classes inheriting just a single property from a schema.org class, but otherwise built entirely from other metadata ontologies can be viewed and registered in the DDE.

We tested the use of the DDE Schema Playground to create customized schema.org-compliant classes that could be used to normalize metadata between multiple types (datasets, clinical trials, publications, etc.) of COVID-19-related resources and applied these schemas towards a searchable resource site (https://outbreak.info). The Outbreak resource schema is available in the DDE schema registry which is also includes schemas from schema.org, Bioschemas, BioLink, the National COVID Cohort Collaborative (N3C), the National Institute of Allergy and Infectious Diseases (NIAID) and more. We hope others will join us in making their open data more interpretable, interoperable, and reusable by adding their schemas to the schema registry.

## Conclusion

We have created a user-friendly browser-based tool which facilitates the application of schema.org towards biomedical research outputs. We demonstrate its use with the creation of the Outbreak.info schema, its adoption into the Bioschemas schema development pipeline, and we encourage others to register and reuse schema.org-compliant schemas.

## Availability and requirements

Project name: Data Discovery Engine Schema Playground

Project home page: https://discovery.biothings.io/schema-playground

Operating system(s): Web-based, Platform independent

Programming language: Python

Other requirements: github account for schema editing

License: Creative Commons Attribution 4.0 International license

Any restrictions to use by non-academics: No

## List of abbreviations

DDE: Data Discovery Engine
FAIR/FAIRness: Findability, Accessibility, Interoperability, Reusability
N3C: National COVID Cohort Collaborative
NIAID: National Institute of Allergy and Infectious Diseases

## Declarations

### Ethics approval and consent to participate

Not Applicable

### Consent for publication

Not Applicable

### Availability of data and materials

Data sharing is not applicable to this article as no datasets were generated or analysed during the current study; however, the Outbreak.info schema generated is available at https://discovery.biothings.io/registry and from gihub at https://github.com/outbreak-info/outbreak.info-resources/tree/master/yaml.

### Authors’ contributions

MC developed the front-end of the DDE Schema Playground with feedback from GT, LDH, and JM. XZ and JX developed the backend of the DDE Schema Playground with feedback from MC. CW and AIS guided the overall development of the DDE Schema Playground. GT developed the Outbreak.info schema with feedback from LDH and JM. GT wrote the manuscript with feedback from LDH, AIS, and CW.

## Acknowledgments

We thank Ben Rush for his suggestions early on in the development of the Outbreak.info schema. We thank the Bioschemas community, especially Nick Juty and Alasdair Gray for their feedback, suggestions, and assistance in improving the DDE and integrating it into the Bioschemas development and maintenance pipeline.

