## Supplemental Figures for "Schema Playground: A tool for authoring, extending, and using metadata schemas to improve FAIRness of biomedical data"

Supplemental Figure 1: Searching and viewing schemas from schema.org and elsewhere

A. Visualizing a json schema file - The schema viewer allows users to visualize json schemas when provided a link to the json schema file

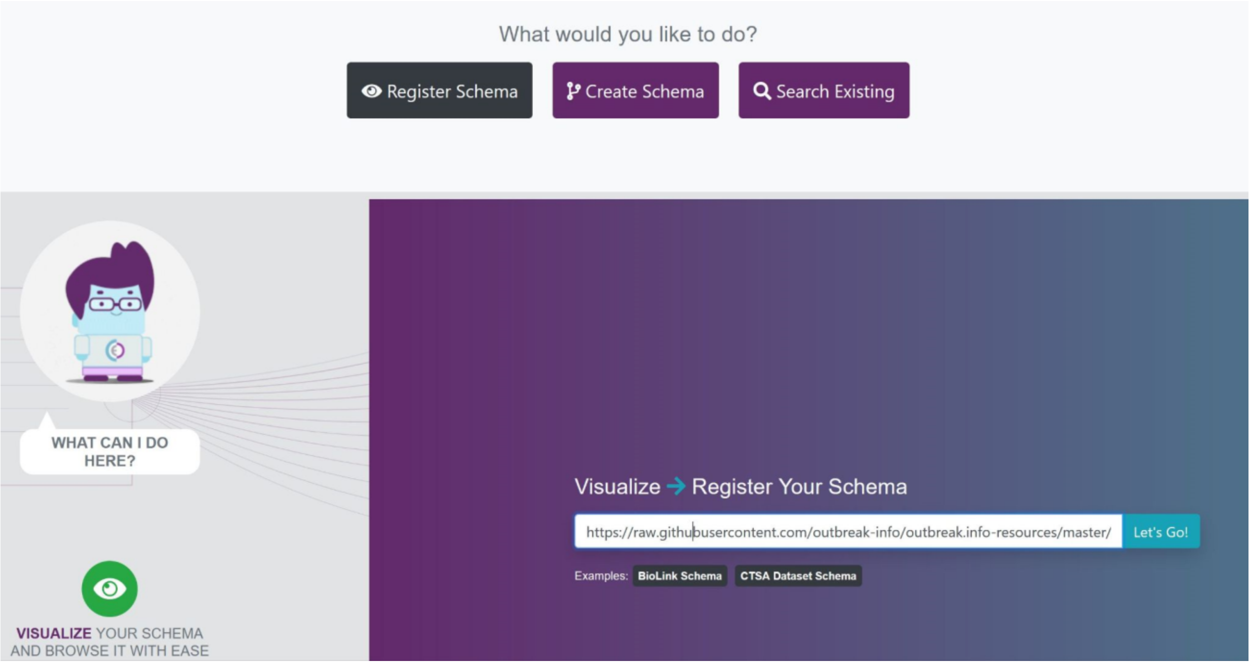


B. Searching/Viewing for a schema in the registry - The Schema registry allows users to browse and view available schema classes so users can find and reuse relevant schemas with ease

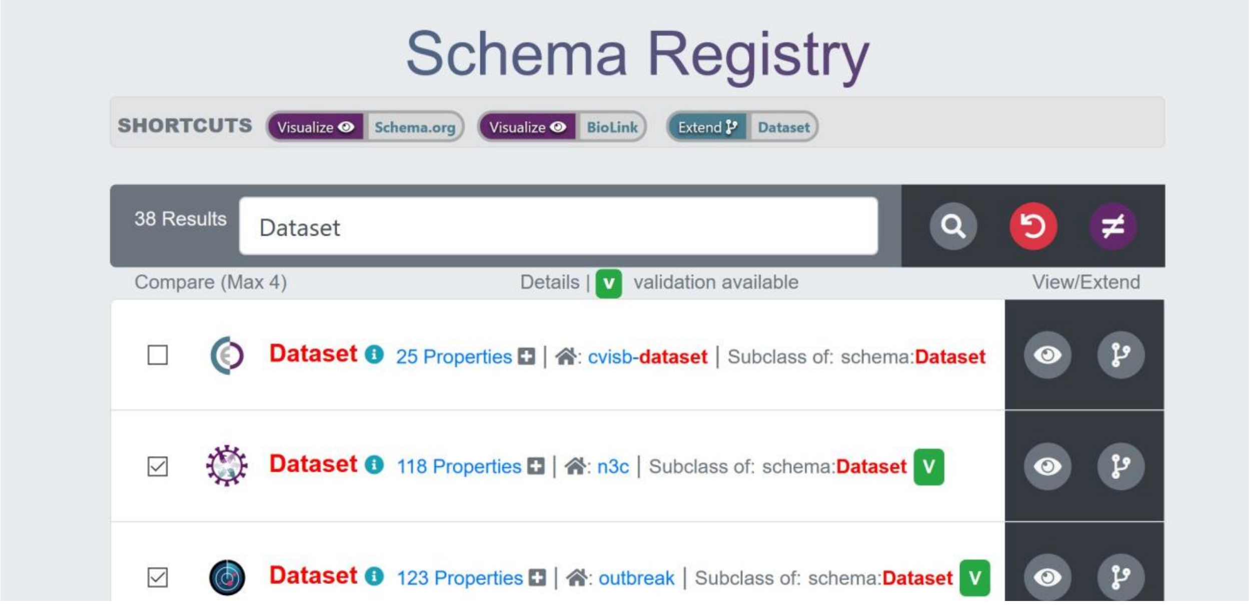

C. Comparing schemas from different sources - If there a similar class exists across multiple schemas, the user can compare the properties that are available (Compare All) or have validation (Compare Used) across the the different schemas

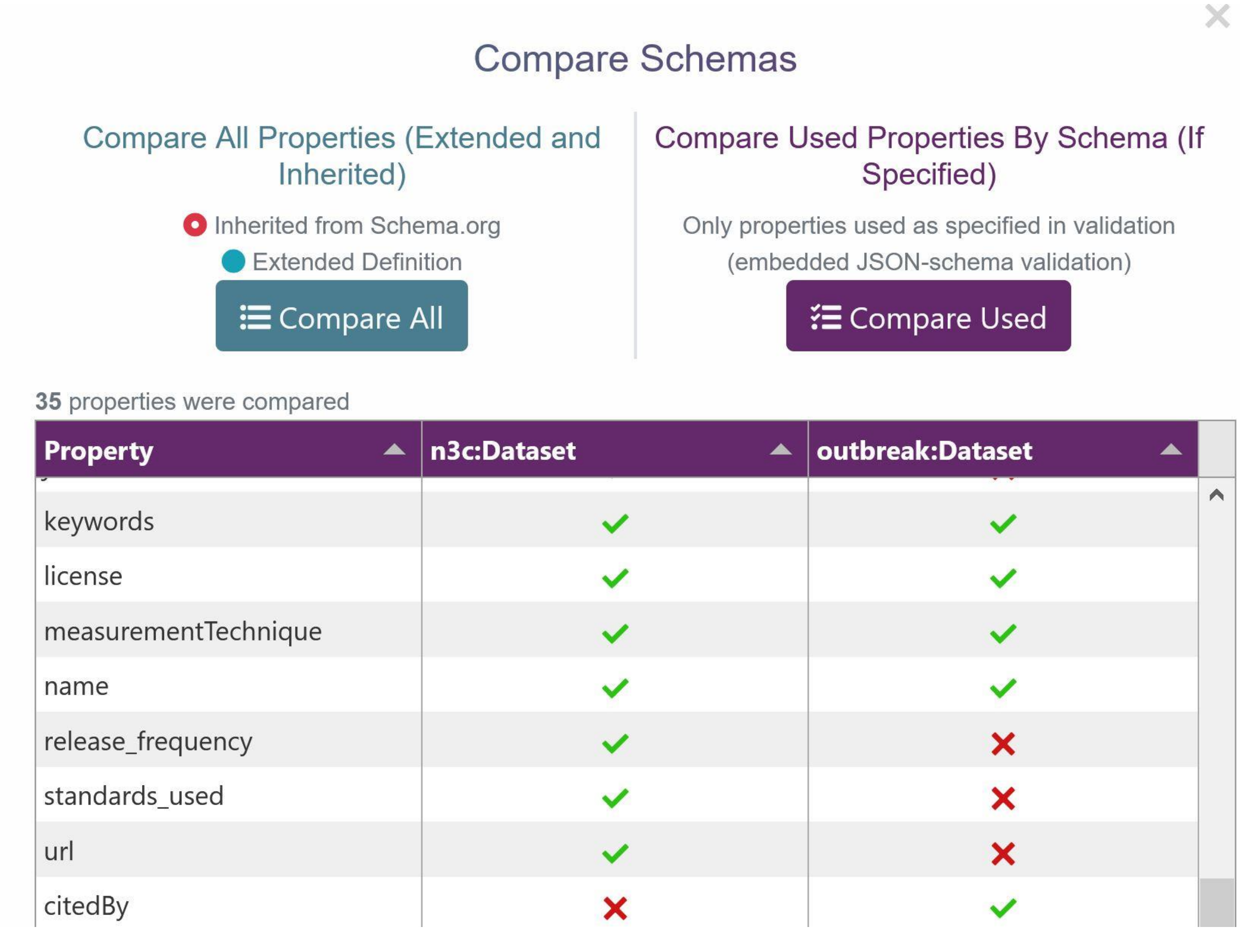


Supplemental Figure 2 - Extending and customizing a pre-existing schema for a particular use

A. Extending/Tailoring a schema from schema.org - If a suitable class for reuse is identified, it can easily be customized by clicking on the 'extend class' icon

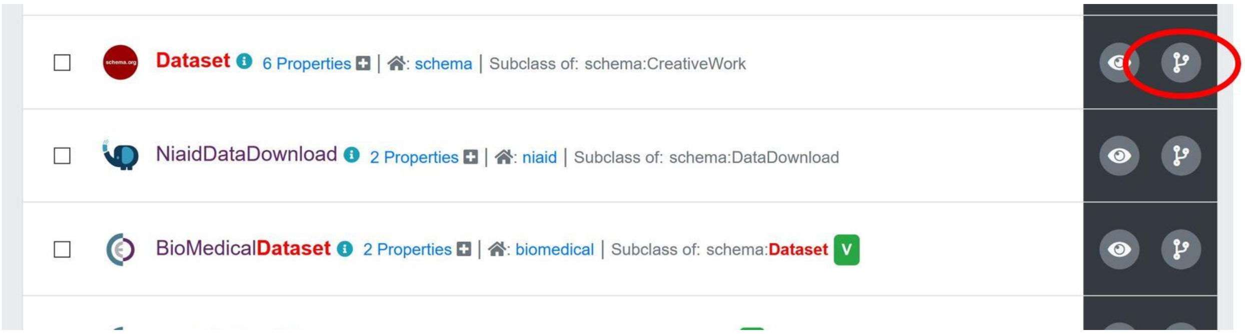

B. Selecting existing properties to inherit/include for validation - To customize a class, the user simply selects properties to be inherited (by clicking on the check icon) or creating new properties (clicking on the + icon)

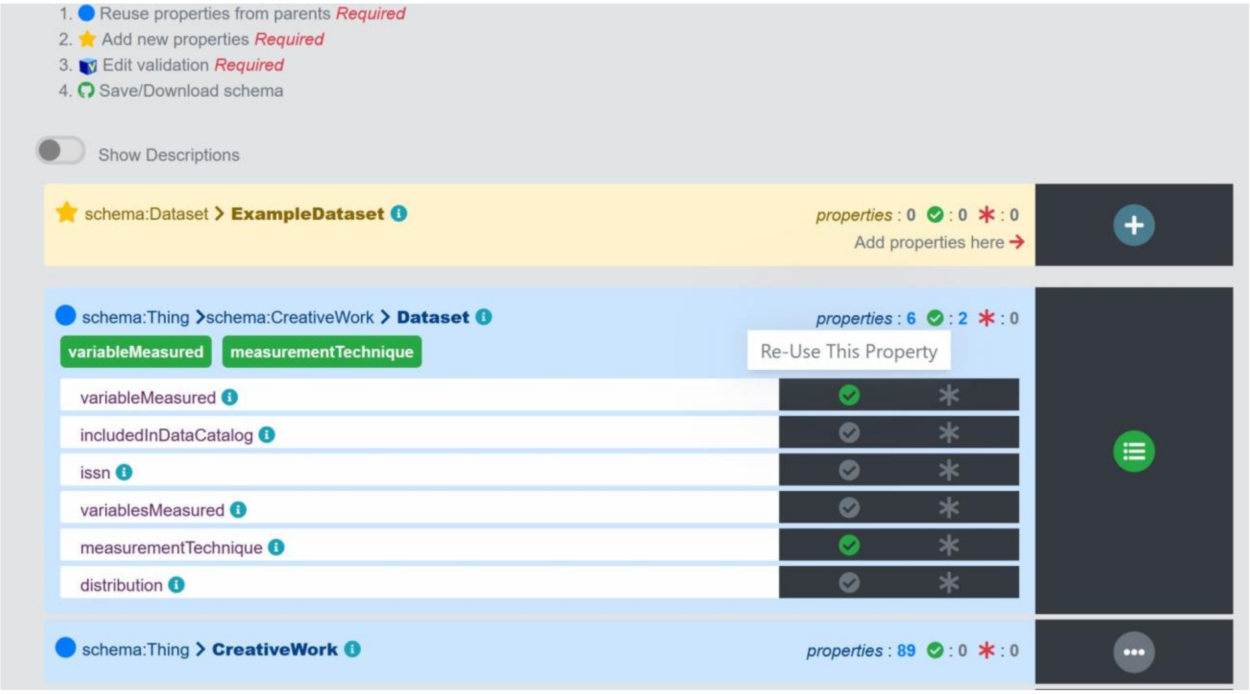


C. Creating new properties - New properties are directed towards conforming to basic schema.org conventions via the property creation guide

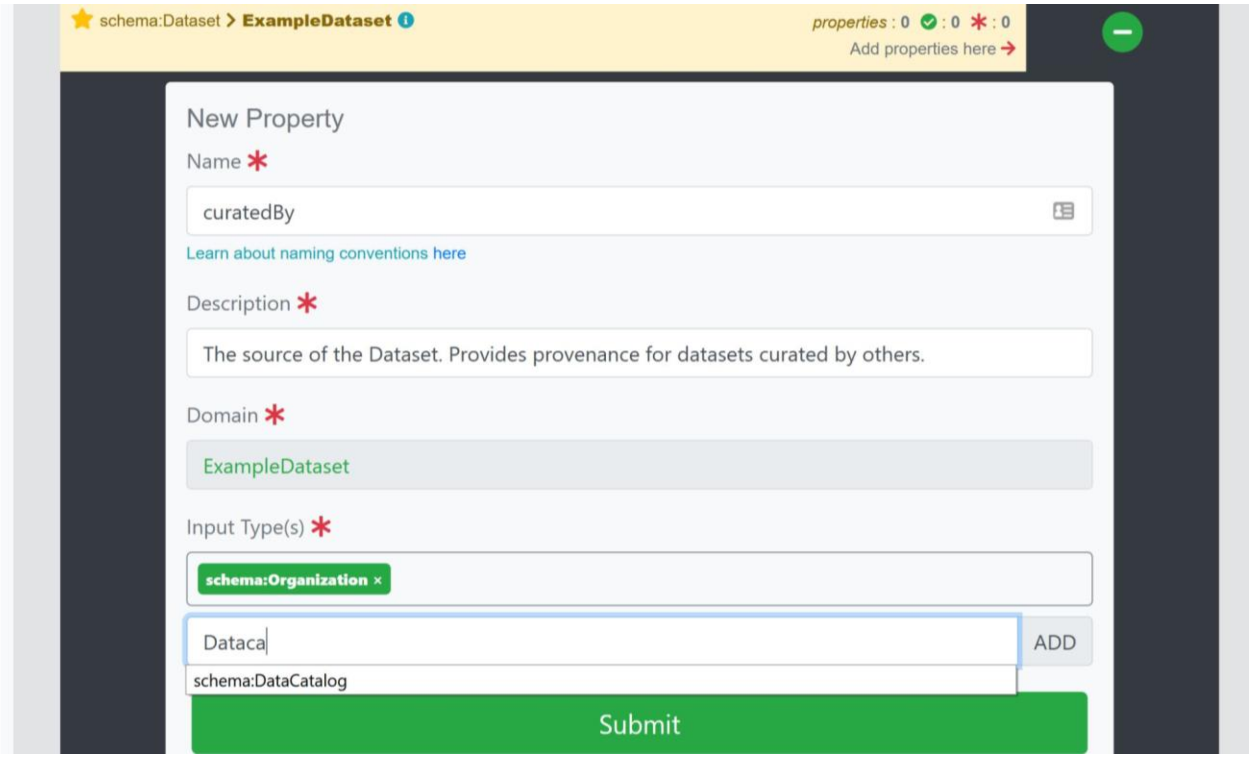

Supplemental Figure 3 - Creating validation for the schema for data quality enforcement

A. Including Marginality - Marginality is enforced in the validation by toggling whether or not a property is required via the * icon

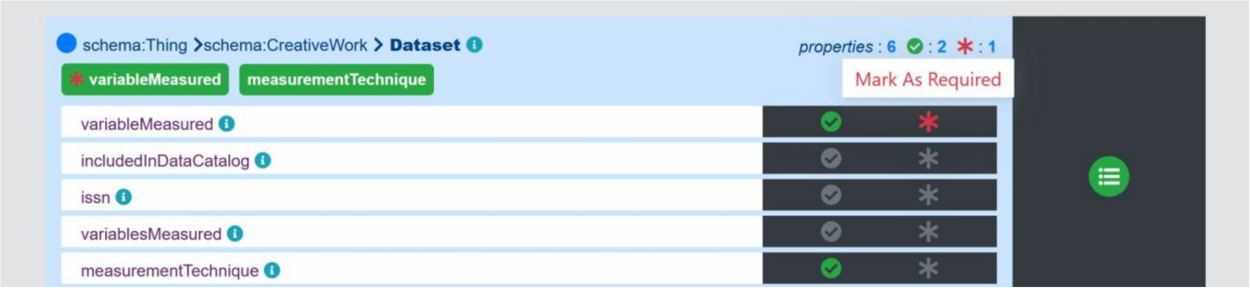


B. Toggling the Validation Editor - Validation can be further customized via the Validation Editor

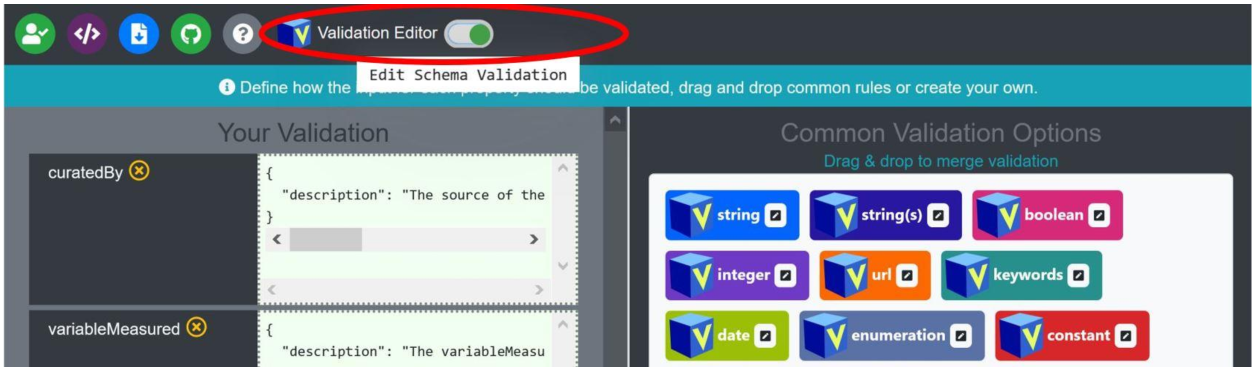


C. Accounting for Cardinality through the validation editor - Cardinality (one/many) is handled via the validation by selecting and editing the validation option. Single options like string or url would be equivalent to a cardinality of one, while an option like string(s) would be equivalent to a cardinality of many (as seen in figure). A cardinality of many for a type other than string can be attained by creating a new validation option (using oneOf or anyOf) or by editing the string(s) option (which uses oneOf). Once the custom validation is saved by clicking on the green ‘save’ button, it will be an option which can be drag/dropped to the appropriate property

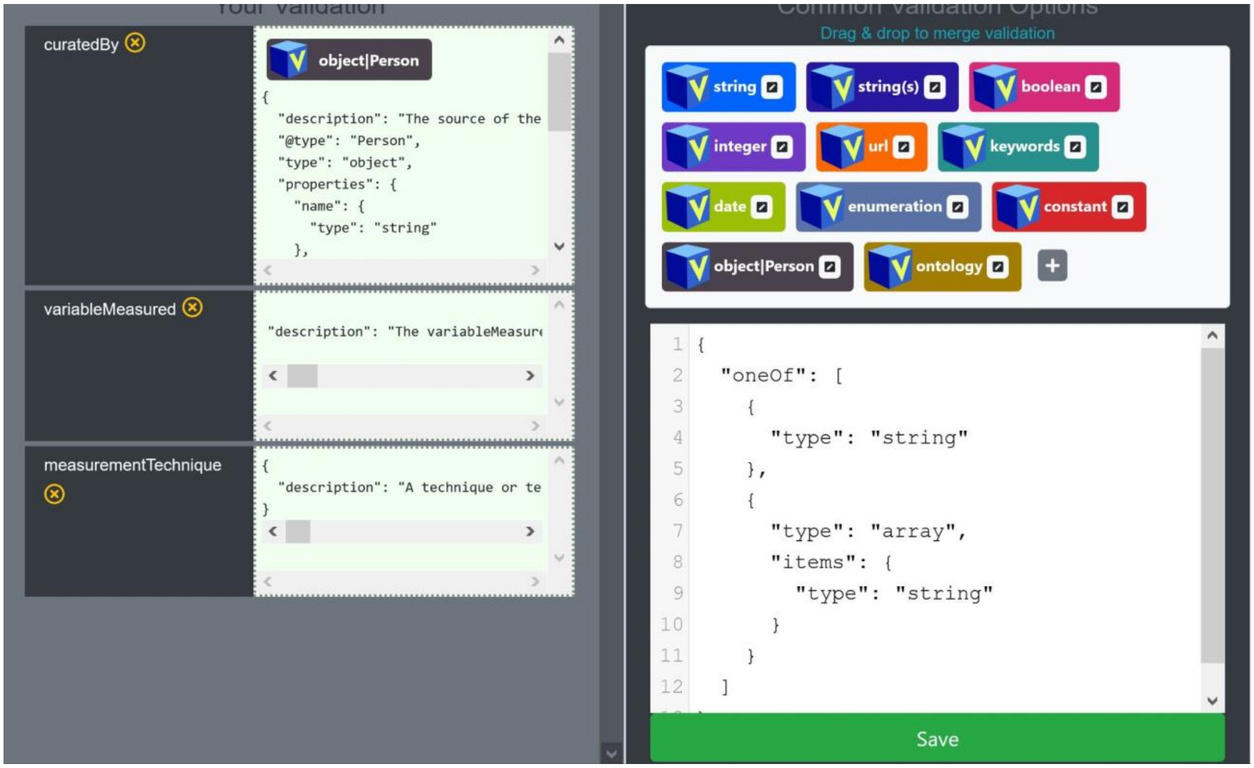


D. Customizing a common validation option - Selecting the (+) option will open a blank editor which will enable the creation of new validation options. To create an organization validation option for the curatedBy property, the content of the object|Person option was copied into a new validation option and edited to replace the “@type” : “Person” with “@type” : “Organization”. Once saved, the option becomes available for drag/drop

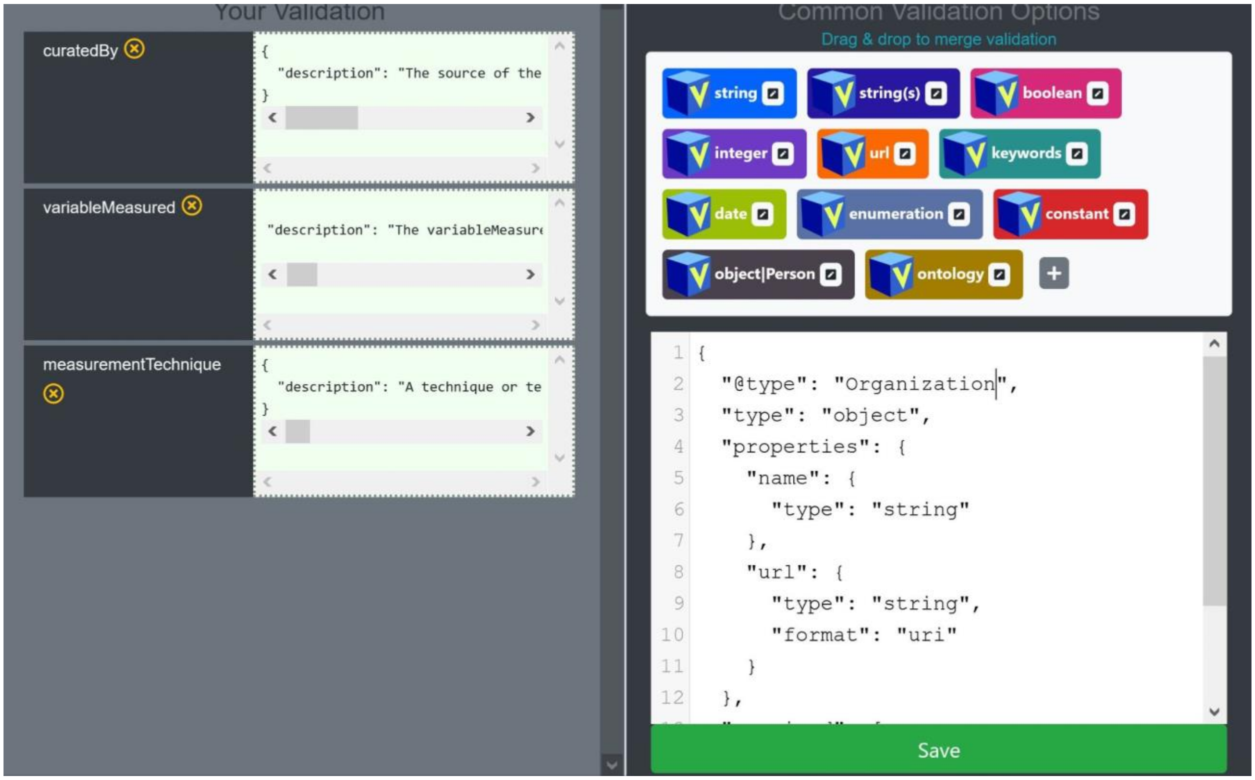


Supplemental Figure 4 - Exporting and saving a schema generated by the Schema Playground editor

A. Exporting/Downloading your schema locally - The schema can be downloaded locally using the download icon

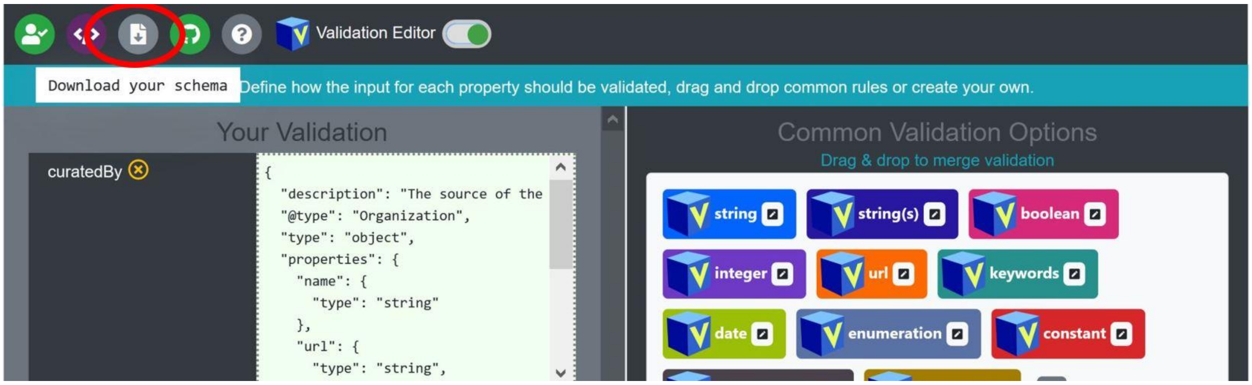


B. Saving your schema to Github - If the user would like to register and share the schema, then it can be saved to a Github repository

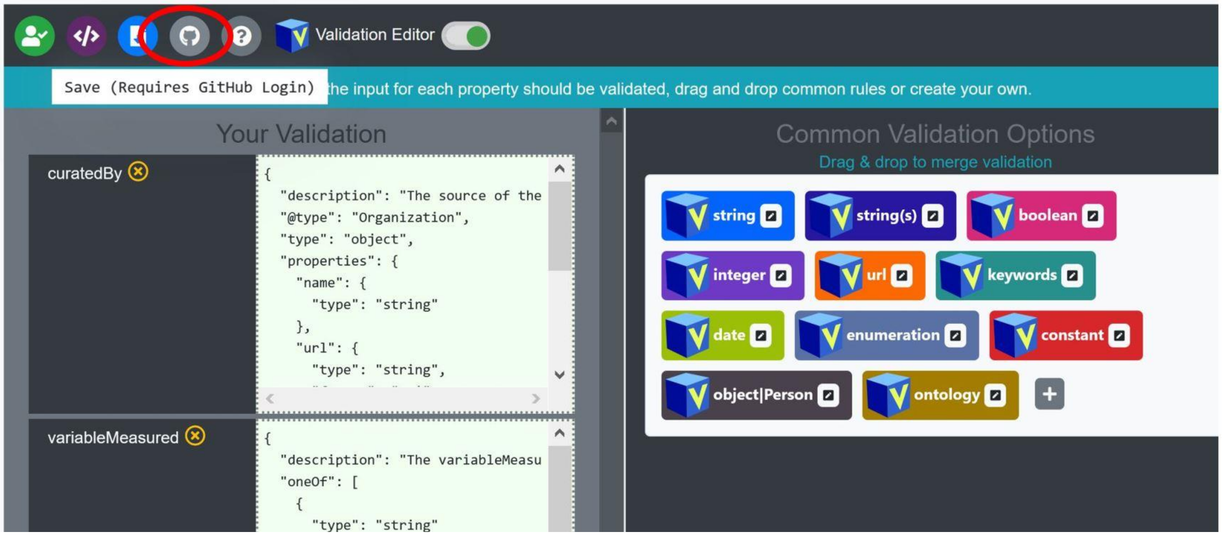


C. Customizing your commit of the schema to Github - When saving to Github, the user can select from their pre-existing repos by clicking on the ‘Get Repos’ button which will populate the available repos dropdown selector. Once a repo is selected, the user can either override an existing file or create a new file. The user can also customize the git commit message/comment

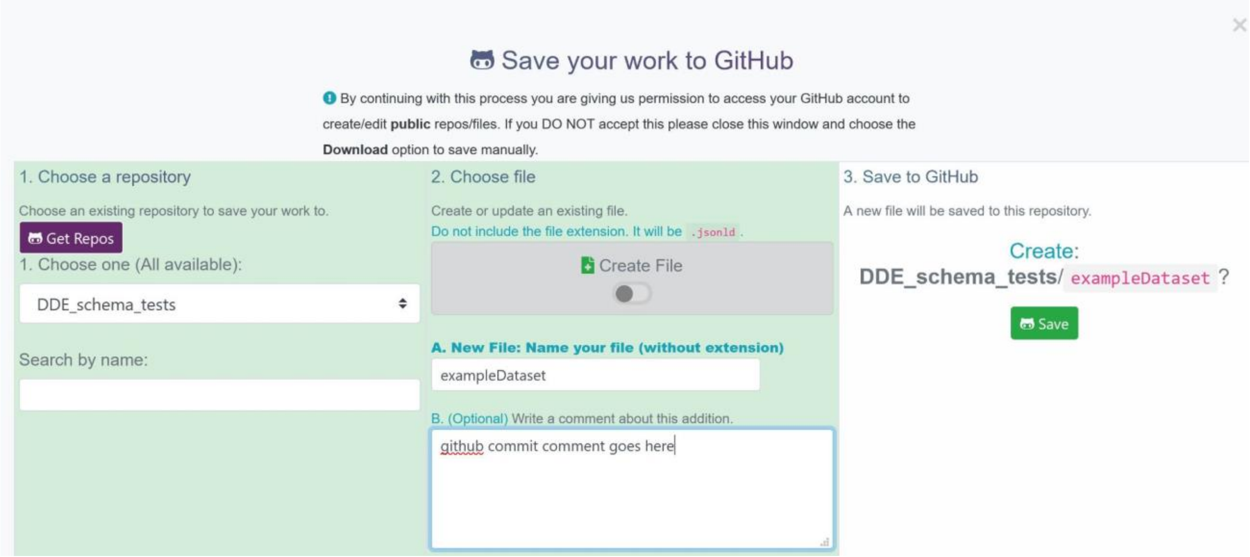
